# Along the Bos Taurus genome, uncover candidate Imprinting Control Regions

**DOI:** 10.1101/2021.12.27.474271

**Authors:** Phillip Wyss, Carol Song, Minou Bina

## Abstract

In mammals, Imprinting Control Regions (ICRs) regulate a subset of genes in a parent-of-origin-specific manner. In both human and mouse, previous studies identified a set of CpG-rich motifs that occurred as clusters in ICRs and germline Differentially Methylated Regions (gDMRs). These motifs consist of the ZFP57 binding site (ZFBS) overlapping a subset of MLL binding units known as MLL morphemes. Furthermore, by creating plots for displaying the density of these overlaps, it became possible to locate known and candidate ICRs in mouse and human genomic DNA. Since genomic imprinting impacts many developmental and key physiological processes, we performed genome-wide analyses to create plots displaying the density of the CpG-rich motifs (ZFBS-morph overlaps) along Bos Taurus chromosomal DNA. We tailored our datasets so that they could be displayed on the UCSC genome browser (the build bosTau8). On the genome browser, we could view the ZFP57 binding sites, the ZFBS-morph overlaps, and peaks in the density-plots in the context of cattle RefSeq Genes, Non-Cow RefSeq Genes, CpG islands, and Single nucleotide polymorphisms (SNPs). Our datasets revealed the correspondence of peaks in plots to known and deduced ICRs in Bos Taurus genomic DNA. We illustrate that by uploading our datasets onto the UCSC genome browser, we could discover candidate ICRs in cattle DNA. In enlarged views, we could pinpoint the genes in the vicinity of candidate ICRs and thus discover potential imprinted genes.

## BACKGROUND

For centuries, breeders have relied on principles of inheritance to obtain animals with desired traits [1]. However, emerging data indicate that assisted reproductive technologies (ART) may induce fetal overgrowth, producing LOS–large offspring syndrome [1–7]. Partly, developmental anomalies arose from altered DNA methylation patterns causing imprinting defects in ICRs, candidate DMRs, or both [4, 5, 7–9]. Furthermore, researchers have observed that Somatic Cell Nuclear Transfer (SCNT) procedures, could epigenetically disturb imprinted gene expression [4].

Overall, genome imprinting is relatively complex and requires orchestrated action of several proteins, including ZFP57, KAP1, and a subset of DNA methyltransferases [10–13]. In ICRs/gDMRs, ZFP57 recognizes its methylated hexameric site [14] and thus plays a central role in the establishment of genomic imprints [10, 14, 15]. ZFP57 family members (KZFPs) are encoded in the hundreds by the genomes of higher vertebrates [16, 17]. Most KZFPs are essential to the recruitment of KAP1 and associated effectors to chromatin to repress transcription [16]. ZFP57 is necessary to maintain the DNA methylation memory at multiple ICRs in mice embryos and embryonic stem cells [10, 14, 15, 18]. In addition to ZFP57 binding sites, the ICRs in mouse often include closely-spaced ZFBS-morph overlaps [19]. These overlaps are composite-DNA-elements that could play dual but antagonistic roles in the regulation of allele-specific gene expression: ZFP57 binding to its methylated sites to maintain allele-specific gene repression; binding of MLL1 or MLL2 to CpG-rich sequences to protect ICRs from methylation to support transcription [19]. MLL1 (or MLL) is the founding member of a protein family with a domain for methylating lysine 4 in histone H3 producing H3K4me3 marks in chromatin [20]. Among the family members, only MLL1/KMT2A and MLL2/KMT2B have the MT domain for binding unmodified CpG-rich DNA [21–23]. Furthermore, through association with several proteins, MLL contributes to the coordinated patterns of gene expression [24, 25].

In both mouse and human DNA, known ICRs/gDMRs encompassed clusters of two or more ZFBS-morph overlaps [26, 27]. Therefore, we wished to investigate whether known bovine ICRs also included these composite-DNA-elements for regulating parent-of-origin-specific expression. To do so, we performed genome-wide analyses of Bos Taurus chromosomal DNA sequences. Firstly, we located ZFP57 binding sites and ZFBS-morph overlaps in the sequences. Subsequently, we created density-plots to pinpoint ICR-positions in Bos Taurus DNA. By uploading our datasets onto the UCSC genome browser, we could obtain snapshots to view peak positions with respect to genomic landmarks. These snapshots uncovered a connection between peaks in plots and the ICRs in bovine imprinting domains including *H19—IGF2*, *KCNQ1*, *PLAGL1*, *IGF2R*, and *PEG3*. Additional snapshots revealed such connection for: the essential ICR in the *GNAS* complex locus; and intergenic ICRs in *MEST*, *INPP5F*, and *MEG8 loci*. Since peaks in plots could locate known ICRs/DMRs in bovine DNA, we anticipate that with our approach one could discover candidate ICRs and novel imprinted genes in cattle DNA.

## RESULTS

For studies of genomic imprinting in cattle, we followed a previous approach applied to mouse and human genomic DNA [26, 27]. Briefly: at the UCSC genome browser, we retrieved the DNA sequences of each cattle chromosome (reported for the build bosTau8). Afterward, we wrote a Perl script to obtain the positions of the ZFP57 binding site and the ZFBS-morph overlaps in bovine DNA. Subsequently, we wrote another script to scan the file containing the positions of the ZFBS-morph overlaps to create density-plots. We tailored this script to count the number of ZFBS-morph overlaps in a sliding window, consisting of 850-bases, while omitting isolated occurrences. The output reported the midpoint for each window as the function of the number of overlap occurrences. We selected the window size by trial and error. Large windows tended to produce false peaks; small windows gave peaks with a spiky appearance. By uploading our datasets onto the UCSC genome browser, we created custom tracks to view the density-plots in the context of landmarks including the positions of genes, transcripts, the CpG islands, and SNPs. Furthermore, by using the browser tools, we could display peak positions in short DNA segments or along an entire chromosome. To evaluate the robustness of our strategy, the following sections offer examples of the positions of ZFP57 binding site, ZFBS-morph-overlaps, and density-peaks in the context of known ICRs or candidate DMRs in bovine DNA. Additionally, we cover an example of how our strategy could be applied to discover novel candidate ICRs and imprinted genes.

### Within the Bos Taurus chromosome 29, density-plots located the ICR of the *H19—IGF2* imprinted domain

In mice, the expression of *H19*, *Igf2*, and *Ins2* is regulated by a single ICR/gDMR positioned upstream of *H19* [12]. *H19* specifies a noncoding RNA gene transcribed from the maternal allele. *Igf2*, and *Ins2* are expressed from the paternal allele and impact fetal growth and body size. As in mice, in cattle the *H19—IGF2* imprinted domain is important to normal growth and fetal development [4, 9, 28, 29]. In cloning studies, deceased newborn calves displayed abnormal expression of *H19* and *IGF2*. In normal surviving adults, the expression of *IGF2* in muscle was highly variable [30]. Furthermore, aberrant methylation of a DMR upstream of *H19* produced abnormal calves and LOS pathogenesis [4, 31]. In *H19* DMR, a study found 33 CpGs in 600 bps [4]. Another study detected a 300-bp DMR at approximately 6 kb upstream of the *H19* promotor [32]. This DMR mapped to a predicted CpG island with a CTCF-binding site corresponding to consensus sequence 5’ -GCGGCCGCGAGGCGGCAGTG-3’ [32].

In the density-plot of Chr29, we noticed a robust peak in a CpG island (CpG45) upstream *H19* (Fig. 1). Peak-position agrees with the reported DMR in cattle DNA [32]. Furthermore, the length of CpG45 is about the same as the CpG-rich DNA (600 bps) selectively methylated in the bovine parental allele [4]. Additionally, under the density-peak, we located 3 predicted CTCF sites (Fig. 1). In our studies, we found predicted CTCF sites using a server at the University of Tennessee Health Science Center [33]. We chose that server because previously it had correctly predicted CTCF binding sites in the ICR of human *H19—IGF2* imprinted domain [34]. The positions of predicted sites agreed with results of ChIPs reporting the association of nuclear proteins (*i.e*. CTCF, RAD21, and SMC3) with chromatin [35].

**Figure 1.**
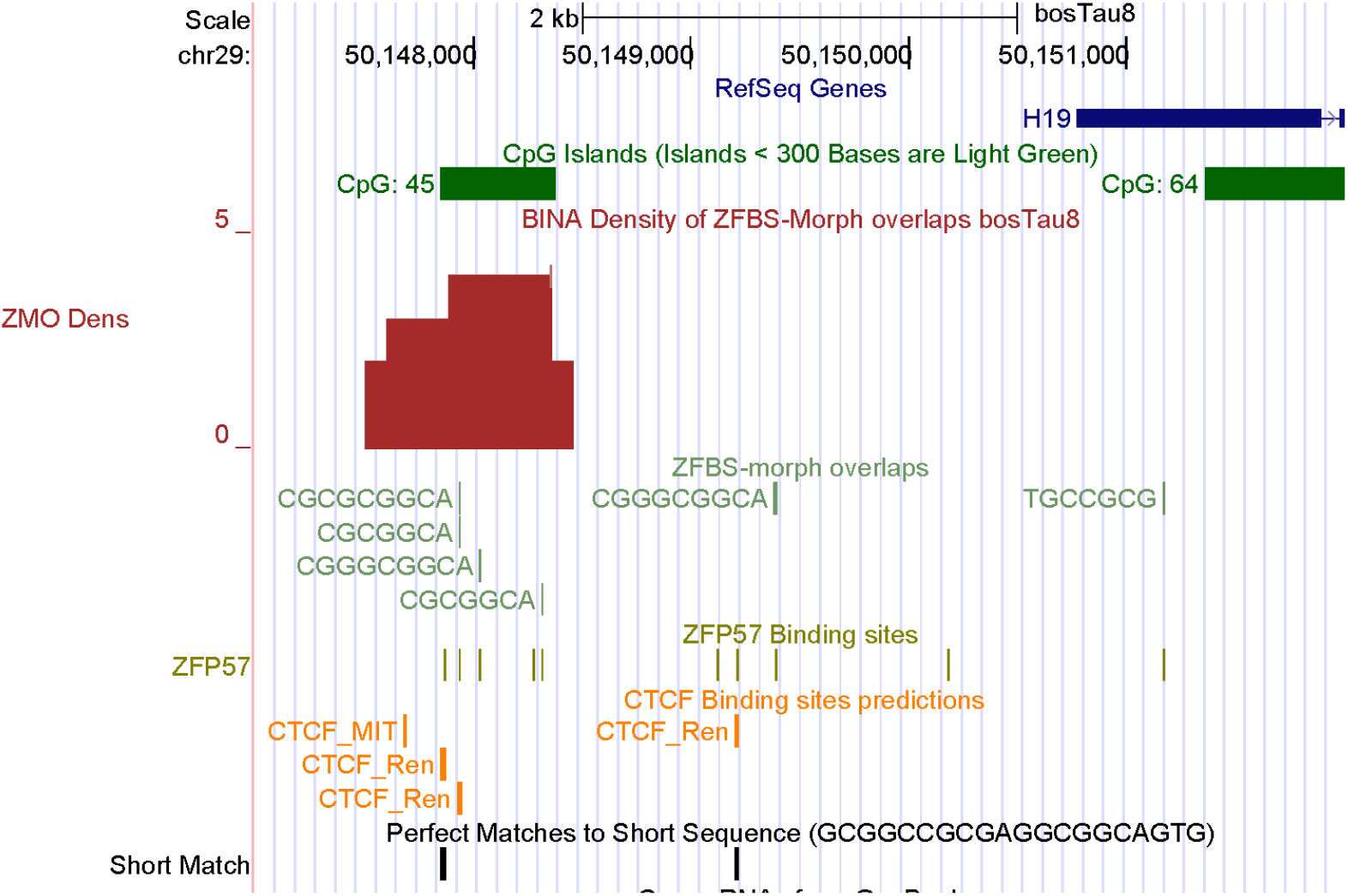
A robust-peak identifying the ICR of the *H19*—*IGF2* imprinted domain in the cattle genome. From top to bottom, tracks display the positions of: cattle RefSeq genes, in pack format; the CpG islands; peaks in density-plot, in full format; ZFBS-morph overlaps, in pack format; ZFP57 binding sites, in dense format. Orange bars mark predicted CTCF binding sites. Short match corresponds to a previously reported consensus CTCF binding sequence. Note: in that plot, the peak is within a previously localized putative DMR encompassing a predicted CTCF-binding sequence.

For studies of cattle DNA, we chose a genomic segment (chr29:50146931-50149221) encompassing the CpG island upstream of *H19* gene in the build BosTau8 at the UCSC genome browser. We submitted the sequence of that DNA segment to the server at the University of Tennessee Health Science Center [33]. In bovine DNA, the server predicted 5 CTCF sites (Fig. 1). One site (CTCF_Ren) corresponds to the consensus sequence (GCGGCCGCGAGGCGGCAGTG) identified previously [32]. Three sites are under the density peak and map to the CpG island that includes the experimentally identified bovine DMR [32]. The 5th site is downstream CpG45 (Fig. 1). Thus, we deduced that our approach correctly located the ICR position in cattle *H19 — IGF2* imprinted domain.

### Within the Bos Taurus chromosome 29, density plots located the ICR of the *KCNQ1* imprinted domain

As in mice [36], the *KCNQ1* imprinted domain in bovine DNA is adjacent to the *H19—IGF2* domain and regulated by an ICR Known as the KvDMR1 [37]. In mice, the KvDMR1 encompasses the *Kcnq1ot1* promotor and regulates imprinted expression of several protein-coding genes [28, 36, 38, 39]. This intragenic ICR is selectively methylated in oocytes but not in sperm [28, 40]. Thus, while *Kcnq1ot1* is transcribed from the paternal allele producing a noncoding RNA, the expression of several genes is repressed in the maternal allele [28]. In mice, targeted deletion of the KvDMR1 caused loss of imprinting and growth deficiency [41]. In cattle, defects in the KvDMR are the most common genomic region affected in LOS pathogenesis [2, 5, 31, 37, 42].

In the density-plots of Bos Taurus DNA, we observed a very robust peak in one of the *KCNQ1* introns (Fig. 2). In order to ensure that this peak is locating the KvDMR1, initially we selected the 5’ and 3’ ends of human *KCNQ1OT1* to perform a BLAT search on the UCSC genome browser. Due to length constraints imposed by BLAT, we could not use the entire human *KCNQ1OT1* as query. Nonetheless, from the BLAT output we could deduce that the peak was in the vicinity of *KCNQ1OT1*. For additional validation, we examined results of a study that localized cattle KvDMR1 using 2 primers to amplify an intragenic DNA.

**Figure 2.**
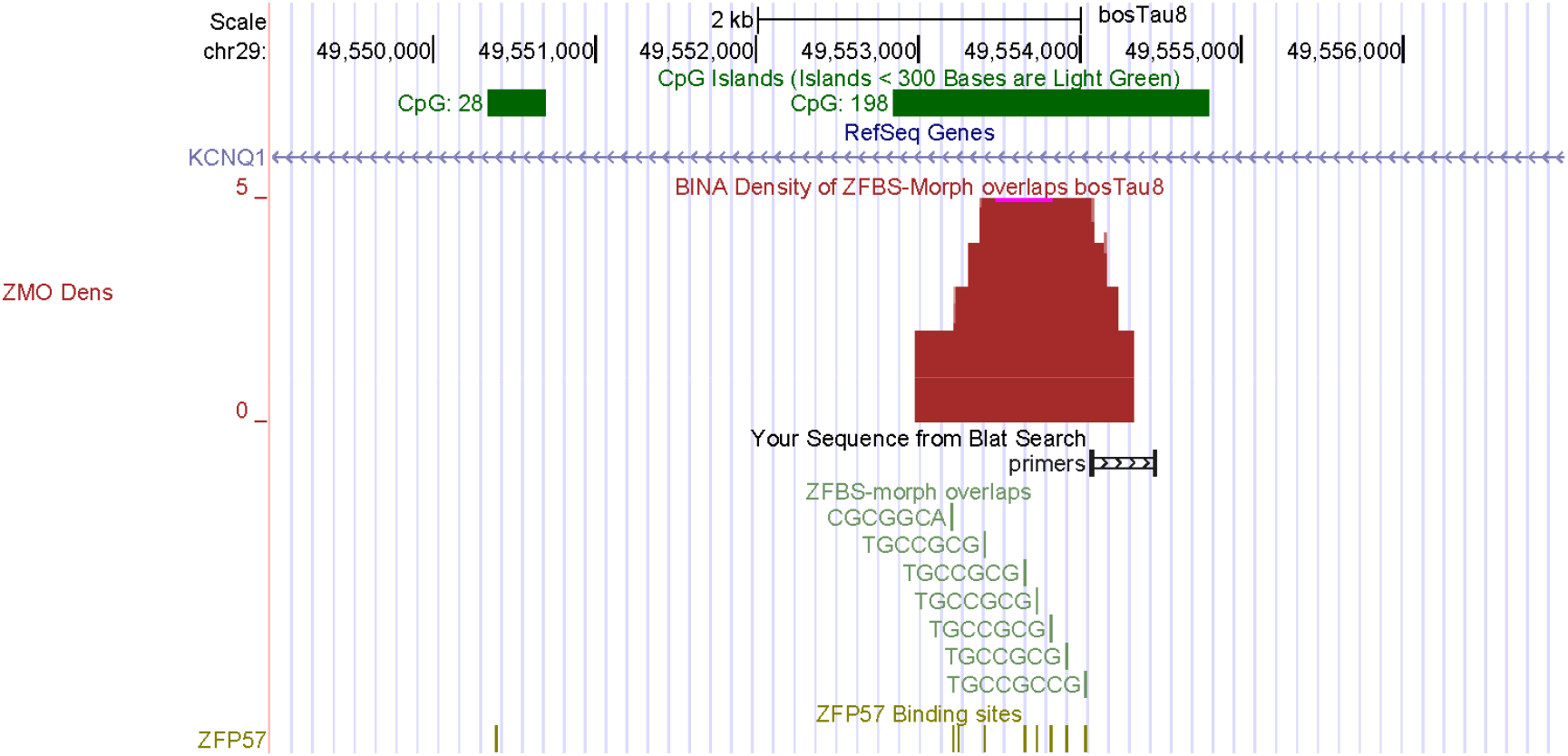
A robust peak pinpointing the KvDMR1 in the cattle genome. Previously, a DNA segment was amplified for locating the KvDMR1 in bovids. The plot shows the position of the two primers used in amplification reaction. These primers are in a CpG island that encompasses the robust peak locating the KvDMR1 in cattle DNA. ZFBS-morph overlaps are shown in packed and ZFP57 binding sites in dense formats.

In that study, statistical analysis showed a significant difference in the methylation level between the two parental alleles –confirming that the amplified DNA corresponded to the KvDMR1 [43]. Therefore, we performed another BLAT search using the sequences of the 2 primers selected for amplifying an intragenic DNA [43]. A close-up view shows that the primers are within a relatively long CpG island that encompasses the robust peak in the density-plots (Fig. 2). Thus, that peak located the central position of the KvDMR in cattle DNA. Notably, while the KvDMR1 in mice encompasses 2 ZFBS-morph overlaps [19], that in cow encompasses 7. Therefore, it seems that ICRs display species-specific differences in the number of ZFBS-morph overlaps they encompass.

### Within the Bos Taurus chromosome 9, density-plots revealed a candidate gDMR/ICRs within the *PLAGL1 locus*

PLAGL1 belongs to a family of zinc finger transcription factors that include PLAG1 and PLAGL2 –reviewed in [44]. One of the *PLAGL1* transcripts (*ZAC1*) encodes a protein that inhibits tumor-cell-proliferation through the induction of cell cycle arrest and apoptosis [44]. In contrast, PLAG1 and PLAGL2 are protooncogenes. Furthermore, *ZAC1* is an intragenic maternally imprinted gene. In both mouse and human genomes, transcription of *ZAC1* originates within an intronic sequence in *PLAGL1*. That intronic region includes another imprinted gene known as *HYMAI* [19, 45, 46]. Both *ZAC1* and *HYMAI* are selectively expressed from the paternal allele. Their ICR/gDMR corresponds to an intragenic CpG island [19, 45, 46]. When the gDMR from human DNA was transferred into mice, this gDMR acted as an ICR and regulated allele-specific expression [47]. In LOS induced by assisted reproduction, imprinting was compromised producing biallelic expression [2, 6].

As previously observed in mice [48], the cattle *PLAGL1* locus encompasses a cluster of several ZFBS-Morph overlaps. In the density-plots, that cluster defines a peak (Fig. 3). However, instead of several *PLAGL1* transcripts, cattle RefSeq Genes displayed only one. We suspected that this transcript corresponded to a previously reported annotation. Specifically, in a panel of maternally imprinted genes, *PLAGL1* was among the loci that acquired methylation marks in an oocyte size-specific manner [49]. In that panel, a putative DMR was localized in a CpG island at the 5’ end of a single transcript referred to as *PLAGL1/ZAC1* [49]. To obtain clues about a complete annotation, we inspected our predicted ICR-position with respect to Non-Cow RefSeq Genes. In the context of human RefSeq Genes, we noticed 2 *PLAGL1* transcripts and a noncoding RNA gene marked as *HYMAI* (Fig. 3). The 5’ end of human *HYMAI* is in the vicinity of an imprinted CpG island [45]. Furthermore, as *ZAC1*, *HYMAI* is expressed from the paternal allele. Therefore: it seems likely that *ZAC1* corresponds to the transcript annotated as *PLAGL1* in cattle RefSeq Genes. Specifically, the genomic organization of mouse *Zac1* is essentially the same as human *ZAC*. Both transcripts originate in a CpG island [46]. In human DNA, *HYMAI* TSS is intragenic [19, 45, 46]. When examined with respect to human RefSeq Genes, the density-peak also is intragenic. Furthermore, in bovine DNA the peak is in a CpG island (Fig. 3). Therefore, we are tempted to conclude that our strategy has pinpointed the reported putative *ZAC1* DMR [49], and to deduce that in bovine *PLAGL1* locus, the putative *ZAC1* DMR is a bona fide ICR.

**Figure 3.**
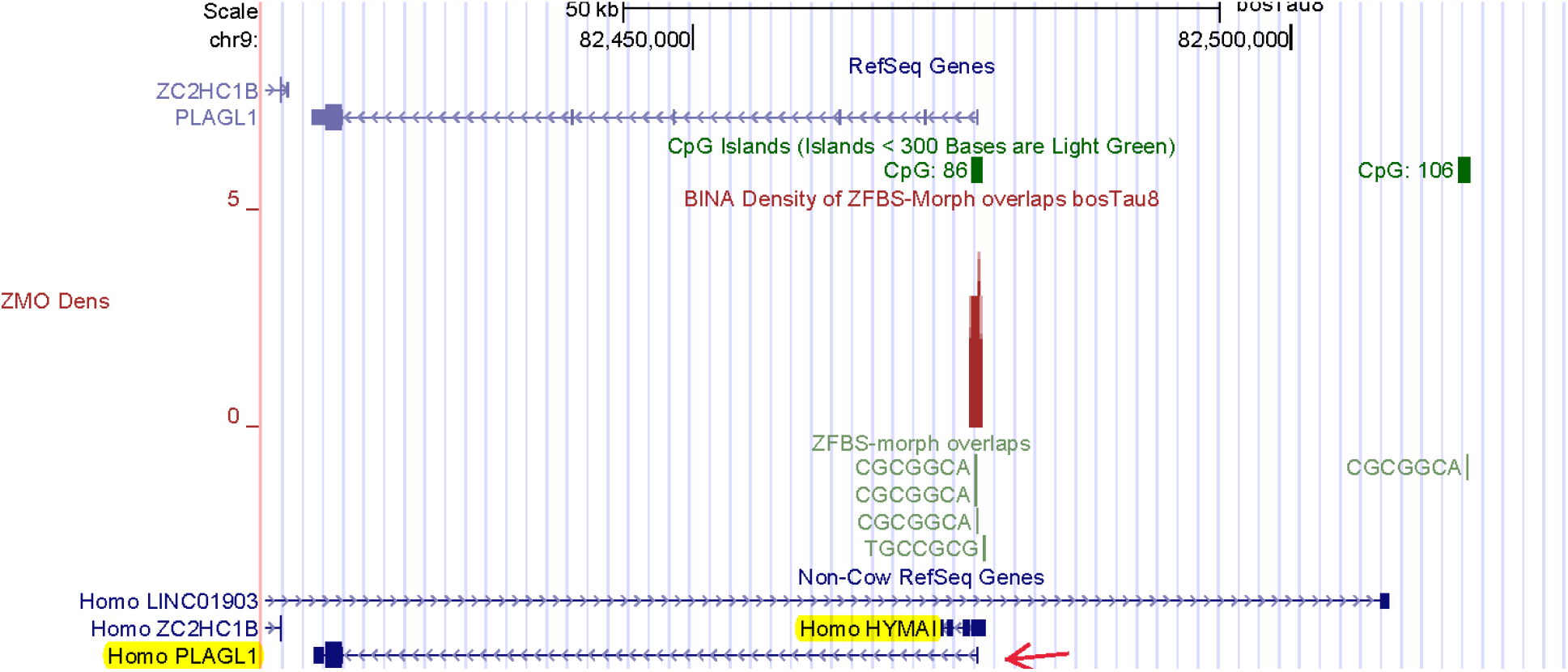
A robust peak predicting the ICR for imprinted expression of the *PLAGL1* transcript known as *ZAC1*. The predicted ICR is at the correct position with respect to two human imprinted transcripts: one transcript corresponds to *HYMAI*; the other to *ZAC1*.

### Within Bos Taurus chromosome 9, density-plots located the ICR for the *IGF2R—AIRN* imprinted domain

IGF2R impacts fetal growth and development. IGF2R also plays key roles in cell differentiation and apoptosis. Its functions include transport of IGF2 into cells and to lysosomes for degradation [50]. In mice, *Igf2r* knockout caused fetal overgrowth and neonatal lethality [51]. The mouse *Igf2r—Airn* imprinted domain includes two differentially methylated CpG islands [52]. DMR1 encompasses the *Igf2r* promotor. The CpG island that encompasses DMR2 is intragenic. In this island is incorporated the promotor of a noncoding RNA gene known as *Air* or *Airn* [53]. DMR2 regulates imprinted expression of *Igf2r* from the maternal and *Airn* from the paternal allele [54]. In Mice, deletion of DMR2 caused biallelic *Igf2r* expression [53]. In cattle, The *IGF2R—AIRN* imprinted domain also includes the DMR2 that regulates imprinted expression. As in mice, DMR2 in cattle is in the second *IGF2R* intron and thus intragenic [4]. As expected, bovine DMR2 is not methylated in sperm and hypermethylated in oocytes [4]. In SCNT experiments, *IGF2R* expression was consistently biallelic –regardless of the source of bovine embryos [4]. Bovine DMR2 is about 2 to 2.7 Kb and located in the second IGF2R intron [4].

While the *IGF2R—AIRN* imprinted domain in Bos Taurus DNA encompasses many CpG islands, one could assume that DMR1 corresponds to the island at the 5’ end of *IGF2R* and DMR2 to the island that maps to the *AIRN* promotor (Fig. 4). In that context, in the density-plots, clearly apparent is a robust peak in the CpG island at the *AIRN* promotor (Fig. 4). Since this peak is in DMR2, we deduced that our approach pinpointed the ICR in the *IGF2R—AIRN* imprinted domain in bovine DNA. Notably, while the gDMR in the mouse genome encompasses 7 ZFBS-Morph overlaps [19], the gDMR in Bos Taurus DNA encompasses 3. Thus, as mentioned above, the number of ZFBS-Morph overlaps in ICRs could be species-specific.

**Figure 4.**
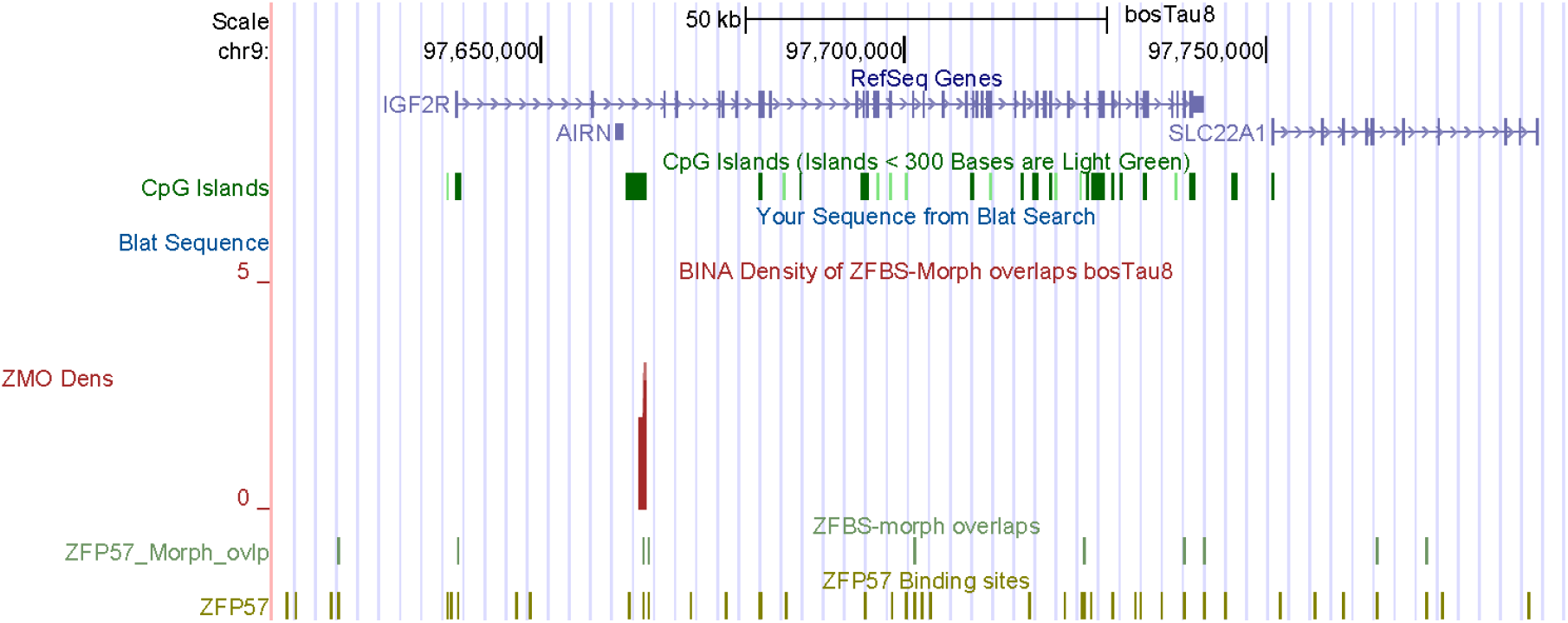
A robust peak defining the ICR of *IGF2R* imprinted domain in cattle genome. As observed for mice, in cattle the ICR in intragenic and maps to the 5’end of *AIRN*.

### Within Bos Taurus chromosome 13, density-plots revealed a candidate for the essential ICR of the complex *GNAS* locus

In both the human and mouse genomes, The *GNAS* locus encompasses multiple DMRs, promotors, and allele-specific transcripts [55]. Since several transcriptional variants are produced from differential exon utilization, they are collectively referred to as *GNAS*. The well-studied *Gnas* locus in mice includes three groups of protein-coding transcripts (*Nesp55*, XL*as*, and G*as*). Although these transcripts share alternative exons, they are regulated by separate promotors [55, 56]. Among the transcripts: *Nesp55* is expressed from the maternal allele; XL*as* from the paternal allele; G*as* from both alleles [56]. *Nesp55* specifies a neuroendocrine secretory protein (known as SCG6). XL*as* and G*as* transcripts are related. Their products function in signal transmission, by G coupled hormone-receptors (GPCRs), and display distinguishable properties [57–60]. The locus also includes an imprinted gene (*Nespas)* transcribed into a noncoding antisense RNA [55].

At the genome browser, we noticed that in contrast to complete annotations reported for mice and humans, cattle RefSeq genes displayed only two transcripts (Fig. 5). From a publication [61], we inferred that the shorter transcript was likely to encode the G*a*s subunit of GPCRs. In cattle, SNPs within that transcript were associated with performance traits [61]. Even though not displayed on the browser, for cattle a previous study identified *GNASXL* as a paternally expressed transcriptional isoform. The study also described a maternally expressed transcript designated as *GNAS* or *NESP55* [62].

**Figure 5.**
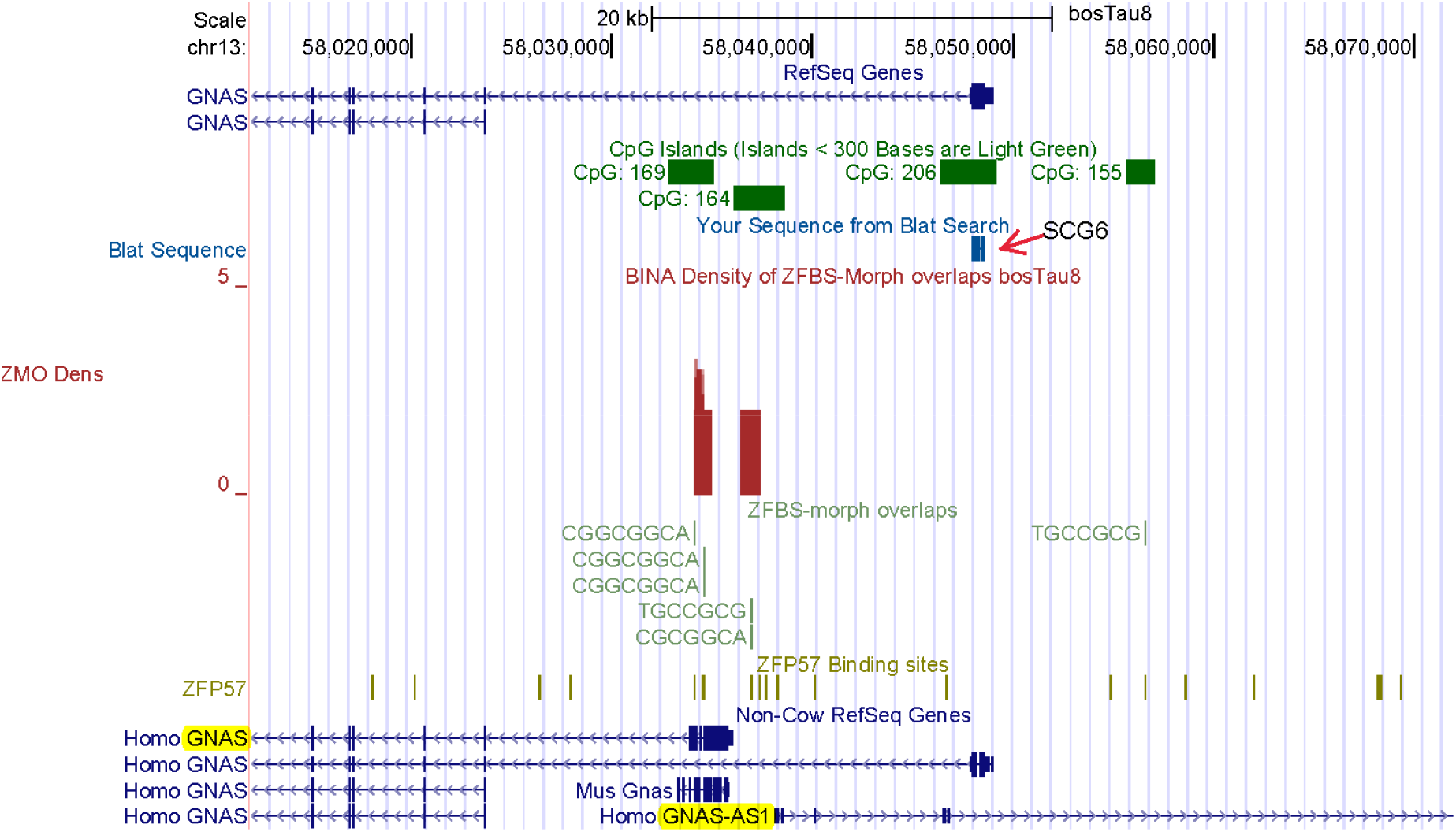
Locating the essential ICR in the *GNAS* locus. The output of a BLAT search identified the longer GNAS transcript as *NESP55* (encoding SCG6). As observed for mice, two peaks define the likely position of the essential ICR of the *GNAS* locus in cattle DNA. The predicted ICR position seems correct with respect to the annotation of Non-Cow RefSeq Genes including human *GNAS-AS1*.

Because the UCSC genome browser’s gene annotations appeared incomplete, we performed a BLAT search to locate *NESP55* in Bos Taurus DNA. For query we chose the amino acid sequence of human SCG6. Based on the output, we deduced that on the browser, the longer *GNAS* transcript corresponded to *NESP55* (Fig. 5). Cattle *NESP55* transcripts are expressed monoallelically in many tissues [62]. Furthermore, in cattle, the *GNAS* locus includes a DMR (a putative ICR) that is hypomethylated in the paternal allele and hypermethylated in the maternal allele [62].

In the density-plots, we observed two clusters of ZFBS-Morph overlaps producing two peaks (Fig. 5). Similarly, a previous study also noticed two clusters of ZFBS-Morph overlaps in the mouse locus [19]. In cattle DNA, both peaks are in the 1^st^ intron of the transcript corresponding to *NESP55* (Fig. 5). Notably, the essential ICR in human locus includes two DMRs: one DMR maps to the first exon of *XLas;* the other is near *GNAS-AS1* TSS [55]. With respect to human RefSeq Genes, peak positions in cattle DNA map to human *XLas* and to a region upstream *GNAS-AS1* (Fig. 5). *In toto*, we could infer that our strategy predicted the genomic position of the essential ICR in the complex *GNAS* locus in the Bos Taurus DNA.

### Within the Bos Taurus chromosome 18, density-plots located a candidate ICR for the *PEG3* imprinted domain

In mice, the *Peg3* imprinted domain encompasses several genes [63]. *Peg3* encodes a relatively large nuclear protein with 12 zinc fingers of C2H2 type [64]. This protein plays a key role in reactions of female mice to her litters [65]. A mutation in *Peg3* caused a striking impairment of maternal behavior. Due to the dearth of maternal care, litters developed poorly and often died [65]. Mechanistically, mutant mothers were deficient in milk ejection –partly due to defective neuronal connectivity, as well as reduced oxytocin neurons in the hypothalamus [65]. In domesticated animals, oxytocin is an indicator of psychological and social well-being [66]. As observed in mice and humans, *PEG3* is expressed from the paternal allele in cattle [9, 67]. In cattle, the *PEG3* domain underwent a global loss of imprinting in fetal overgrowth syndrome induced by assisted reproduction [2].

Sequence analyses have predicted that in Bos Taurus, the *PEG3* imprinted domain encompasses many CpG islands [67]. Consistent with this prediction, at the genome browser we observed several CpG islands in bovine DNA (Fig. 6). In cattle, the island encompassing the *PEG3* promotor was the only area that showed DMR status [67]. Notably, in the density-plots we observed a peak in that DMR (Fig. 6). Since the DMR was methylated in an allele-specific manner [67], we could deduce that the peak pinpointed the ICR in cattle *PEG3* imprinted domain. Furthermore, earlier studies of mice revealed that clusters of ZFBS-morph overlaps mapped to functionally important landmarks –including DNase I hypersensitive sites and repressive H3K9me3 marks [19]. In mouse, these clusters were in the 1^st^ *Peg3* intron. Likewise, for cattle we observe an intragenic peak. Therefore, our strategy pinpointed the central portion of the ICR in cattle *PEG3* imprinted domain (Fig. 6).

**Figure 6.**
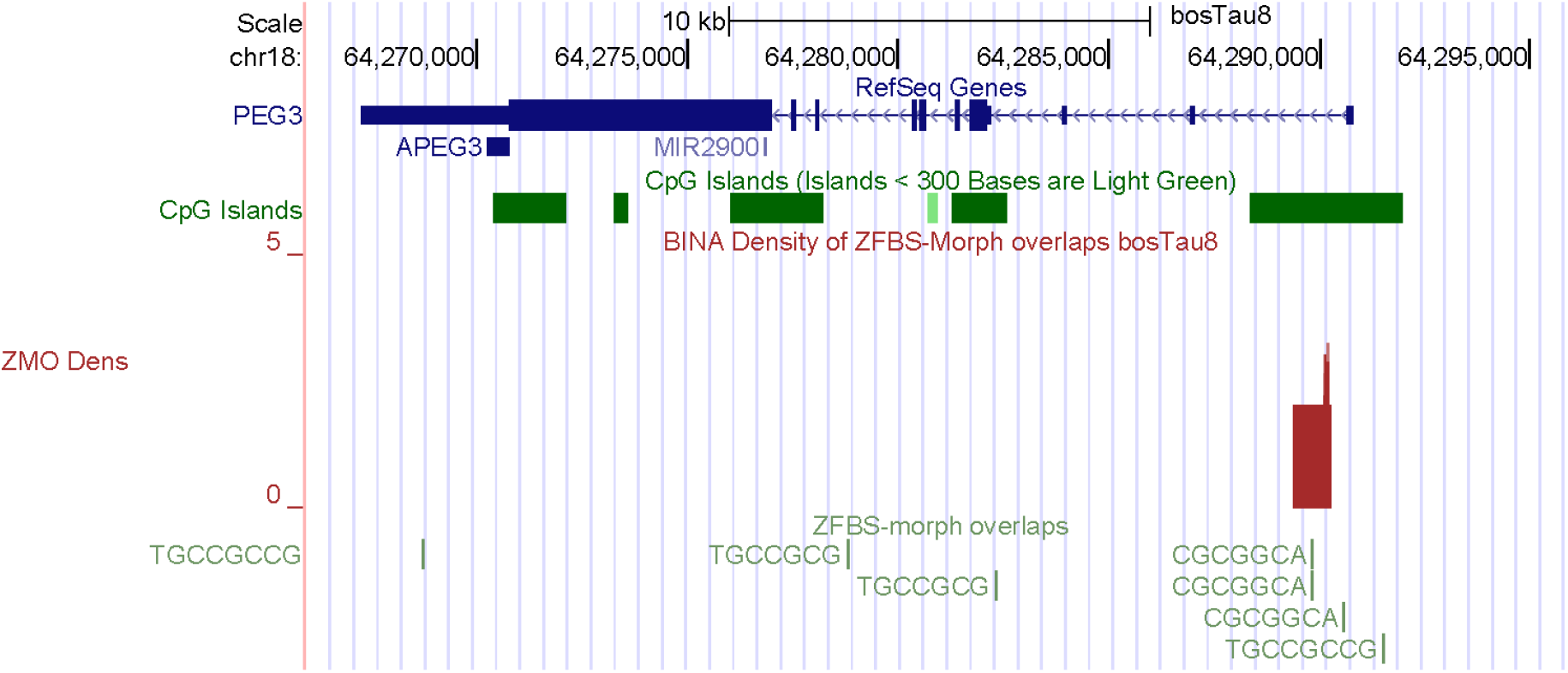
A robust peak locating the ICR of *PEG3* imprinted domain. As detailed in the text, this peak is within the CpG island that is methylated in a parent-of-origin-specific manner.

### Within Bos Taurus chromosome 4, density-plots located the intragenic ICR regulating allele-specific expression of *MEST* isoform

MEST is a member of the α/β-hydrolase fold family of enzymes [68]. In mice, the TSS of a paternally expressed transcript originates from the 1^st^ *Mest* intron [68, 69]. This transcript is also known as *Peg1*.An intragenic ICR regulates its expression selectively from the paternal allele. In cattle, among 8 investigated genes, only *MEST* showed differential expression in day 21 parthenogenetic embryos [70]. In mice, the product of *Peg1* impacts maternal behavior [71]. This conclusion was reached from studies of *Peg1* deficient females. Even though these females displayed a normal investigative behavior, they lacked appropriate maternal response [71]. Consequently, their newborn litters had a low survival rate [71]. Furthermore, *Peg1* deficient females displayed abnormal maternal activities including impaired placentophagia.

In bovine DNA, methylation imprints are established in an oocyte size-specific manner [49]. In a panel of maternally imprinted genes, putative DMRs were localized in several imprinted loci including *MEST*. CpG analyses deduced that the putative *Peg1/MEST* DMR maps to a CpG island [49]. Consistent with this prediction, density-plots revealed a peak in a CpG island in the *MEST* locus (Fig. 7). Since on the browser we observed a single *MEST* transcript, we inspected peak-position with respect to Non-Cow RefSeq Genes. In the context of human RefSeq Genes, the peak is upstream of *MESTIT1* and near one of the *MEST* short isoforms. As observed for *MEST*, *MESTIT1* is expressed from the paternal allele [72]. Thus, the density-plots correctly pinpointed the ICR regulating the expression of *PEG1/MEST* transcript in bovine DNA. Furthermore, the plots demonstrated the correspondence of this predicted ICR to the reported imprinted DMR in cattle DNA [49].

**Figure 7.**
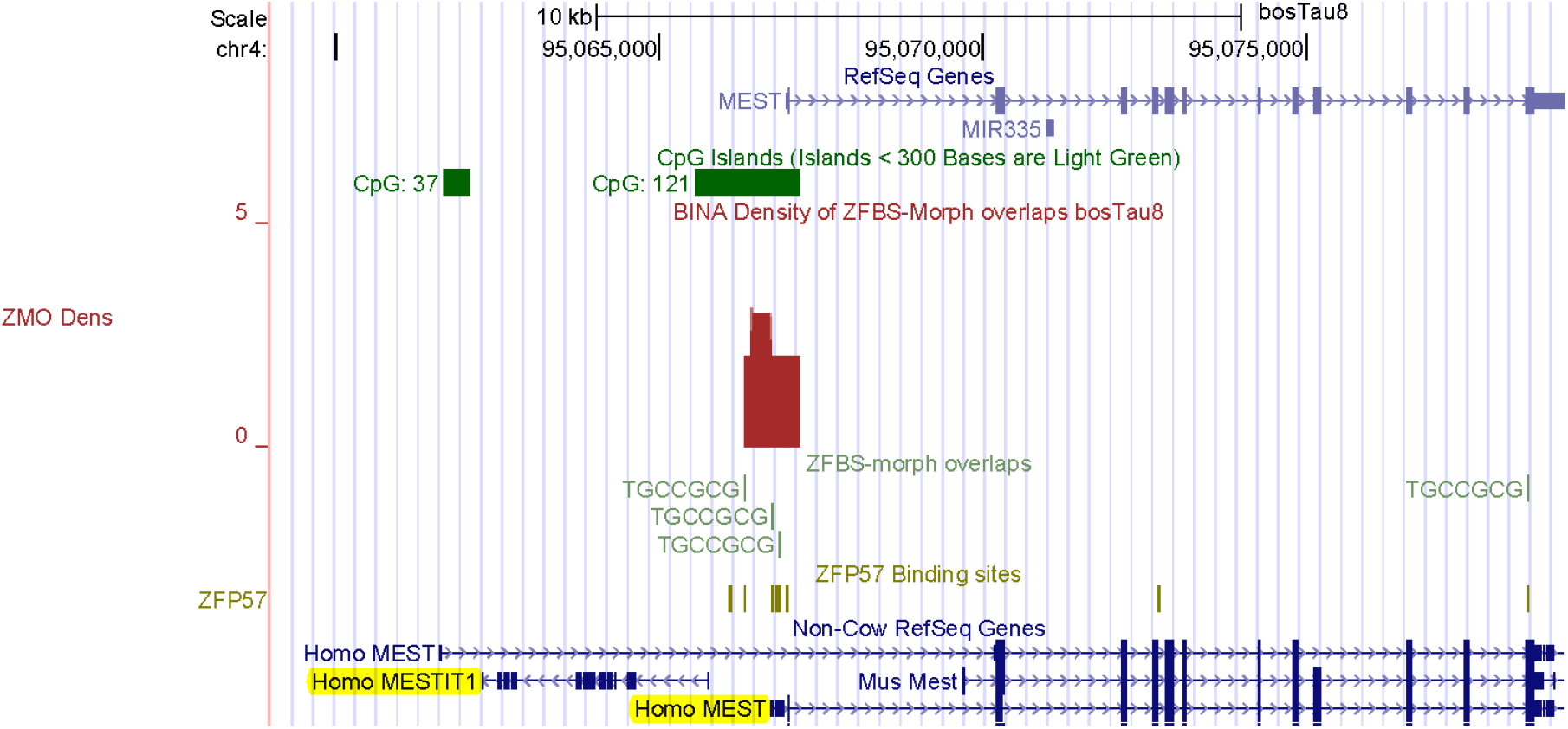
A robust-density-peak locating the ICR for imprinted expression of the *MEST* variant in cattle DNA. This location is presented in the context of Non-Cow RefSeq Genes for human *MESTIT1* and *MEST* transcriptional variant.

### Within Bos Taurus chromosome 13, an intragenic peak located the ICR regulating allele-specific expression of *NNAT*

The neuronatin gene (*NNAT*) is expressed during postnatal brain development [[73]. In mice, *Nnat* is located within the single intron of *Blcap* [74]. Also known as *Peg5, Nnat* is an imprinted gene. *Blcap* is expressed biallelically [74]. From *Nnat* are produced several alternatively spliced isoforms [74]. In human, the CpG island in *NNAT* promotor was differentially methylated in all examined tissues [75]. Therefore, that island corresponds to the ICR that regulate *NNAT* expression [75].

As in humans and mice [74–76], in cattle two *NNAT* transcripts are expressed from the paternal allele [77]. In plots of cattle DNA, we noticed a CpG island near the *BLCAP* TSS. Another island (CpG59) is intragenic (Fig. 8). In the density-plots, we did not find a peak in the island near *BLCAP*, a finding that is consistent with *BLCAP* biallelic expression. In *BLCAP* intron, we noticed 2 peaks. One of the peaks is more robust. It is located in CpG59 and encompasses both *NNAT* TSS and promotor (Fig. 8). This peak is the likely position of the ICR that regulates imprinted *NNAT* expression in bovine DNA Fig. 8). Overall, robustness of peaks depends on the number of ZFBS-morph overlaps they encompass. Peaks that cover 2 ZFBS-morph overlaps could be true or false positive [26]. Peaks that cover 3 are more robust.

**Figure 8.**
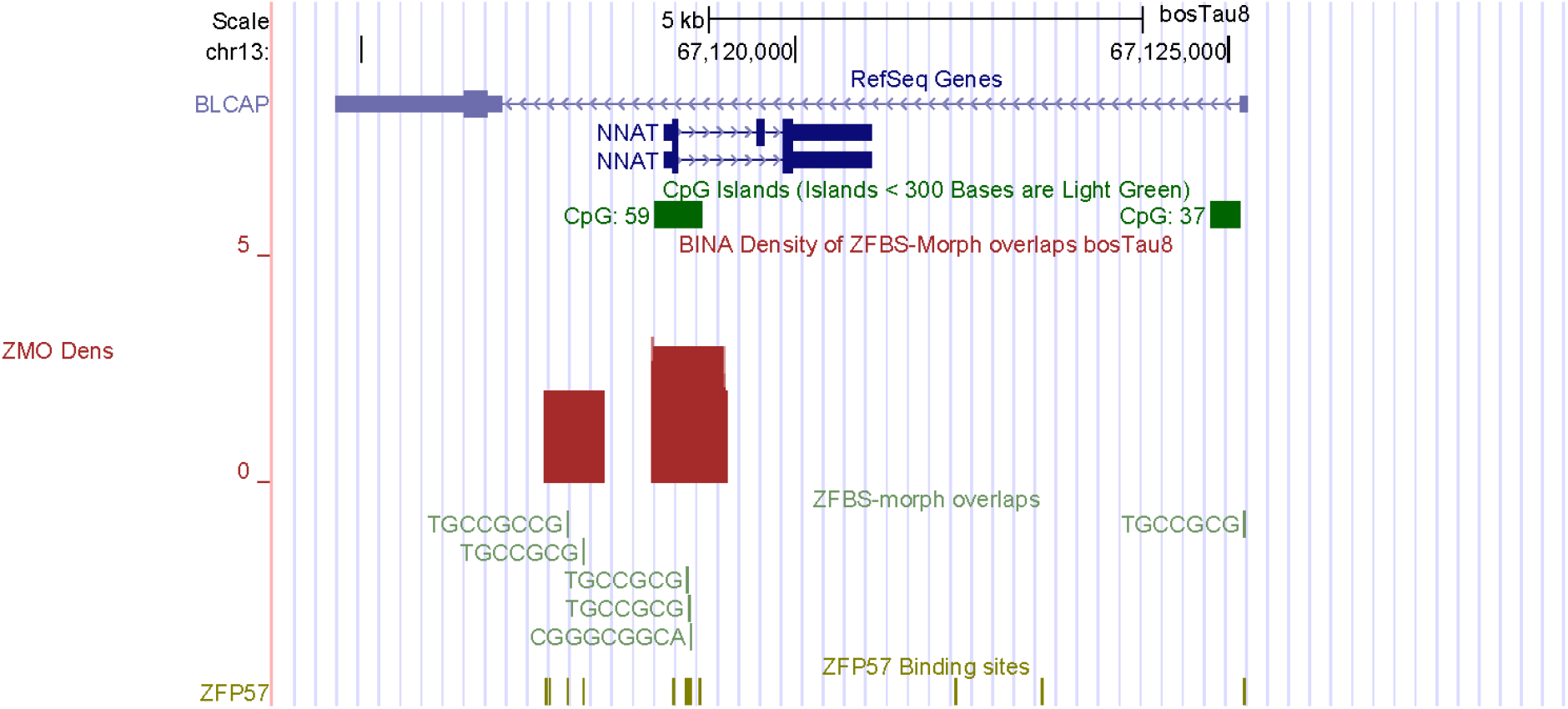
A robust-density-peak locating a candidate ICR for imprinted *NNAT* expression in cattle DNA. This peak is in CpG59 and maps to a known cattle DMR.

### Within Bos Taurus chromosome 21, a peak predicted a candidate ICR for regulating expression of *MEG8*

In cattle, the *DLK1-DIO3* imprinted cluster includes a maternally-expressed gene (*MEG8*) transcribed into a noncoding RNA [78]. In an 8-wk-old animal, *MEG8* was preferentially expressed in skeletal muscle [78]. In Angus calves, maternal diet during pregnancy impacted *MEG8* expression in longissimus dorsi muscle [79]. In adult cattle, *MEG8* was expressed in several tissues –including heart, liver, spleen, lung, kidney, brain, subcutaneous fat and skeletal muscle [80]. In heterozygous cattle, *MEG8* was expressed from only one of the two parental alleles. In the density-plots, we observed a robust peak predicting a candidate ICR for allele-specific expression of cattle *MEG8*. This peak is intragenic and maps to a CpG island (Fig. 9).

**Figure 9.**
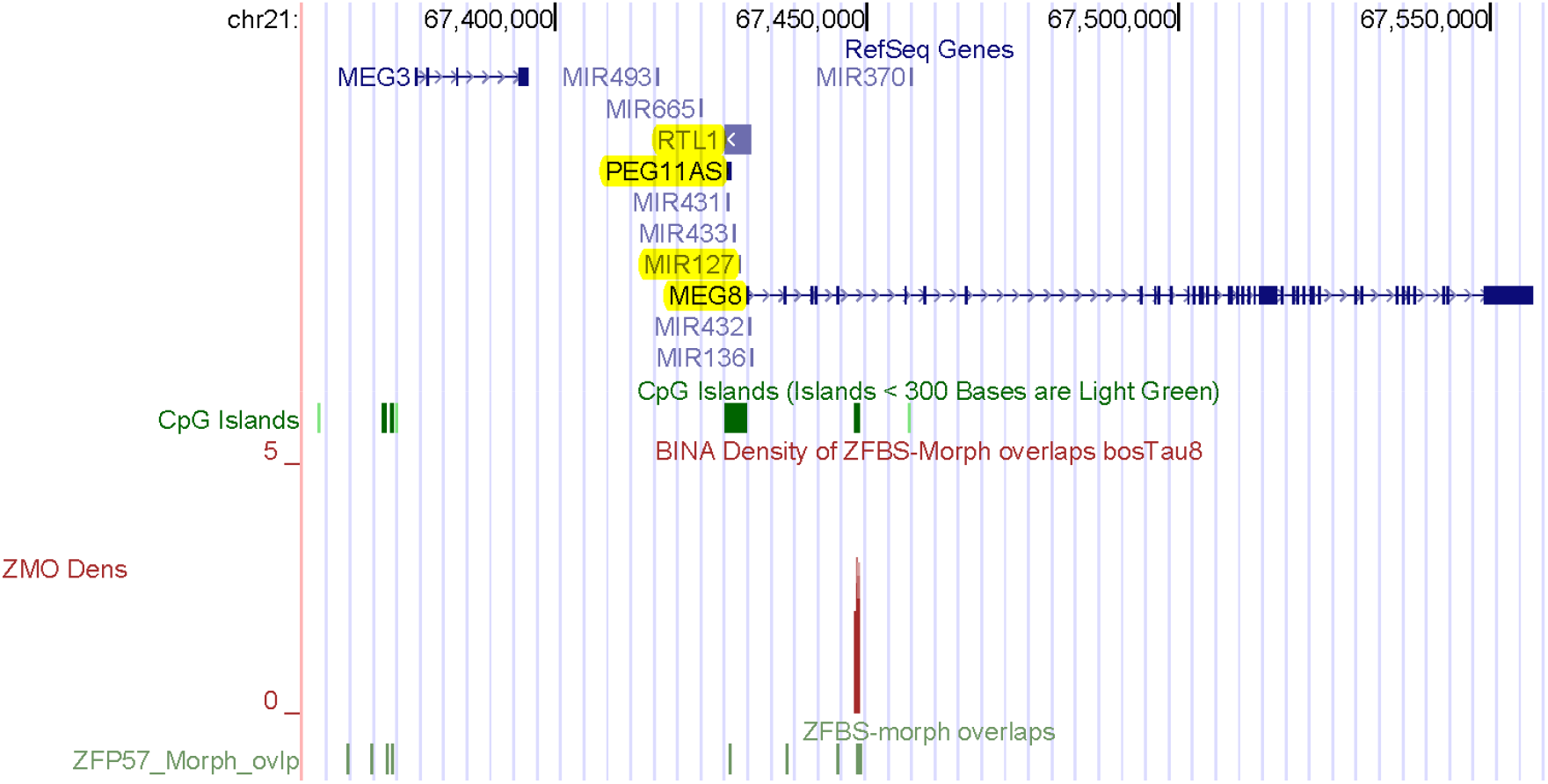
A robust peak predicting the ICR for imprinted expression of *MEG8*. This peak is intragenic and maps to a CpG island.

### Within Bos Taurus chromosome 26, a peak predicted a candidate ICR for imprinted *INPP5F_V2* expression

INPP5F is an inositol 4-phosphatase that functions in the endocytic pathway [81]. In mice, *Inpp5f_v2* was identified as an imprinted gene in the brain [82]. *Inpp5f_v2* is a variant of *Inpp5f*. It is a retrogene with a unique alternative first exon. *Inpp5f_v2* transcription originates in one of the *Inpp5f* introns. While *Inpp5f* is biallelically expressed, *Inpp5f_v2* is transcribed from the paternal allele [82]. Furthermore, mouse *Inpp5f* and human *INPP5F* share sequence similarity. In mouse *Inpp5f* locus, CpG analyses identified two CpG islands [82]. CpG1 is near the 5’ end of *Inpp5f*, CpG2 is at 5’ end of *Inpp5f_v2*. Bisulfite sequencing of the CpG1 island showed that both alleles were hypomethylated, as would be expected for a nonimprinted transcriptionally active gene. In the brain, CpG2 was methylated on the maternal allele, but not on the paternal allele. Thus, the intragenic island regulates imprinted *Inpp5f_v2* expression [82]. Similarly, the promotor of human *INPPF5_V2* transcript is embedded within a maternally methylated DMR [83].

Literature surveys we could not find reports concerning cattle *INPPF5_V2*. Furthermore, the genome browser did not include any annotation for the *INPPF5* locus in cattle DNA. The only annotated gene corresponded to *MCMBP* (Fig. 10). However, we were able to locate *INPP5F* transcripts in the context of Non-Cow RefSeq Genes. With respect to human genes, *INPPF5* is upstream of *MCMBP* –as observed in human and mouse genomic DNA. Since we were in correct genomic area in cattle DNA, we could ask whether the density-plots included a candidate ICR for allele-specific expression. In plots, we noticed a robust peak in a CpG island (CpG55) in cattle DNA. With respect to human transcripts, this peak maps to the 5’ end of *INPPF5_V2*. Therefore, CpG55 is the likely ICR position in *INPPF5* locus in cattle DNA. As first described for mice, this ICR is intragenic [82]. Thus, our strategy predicted a candidate ICR for imprinted bovine *INPPF5_V2* expression from bovine DNA (Fig. 10).

**Figure 10.**
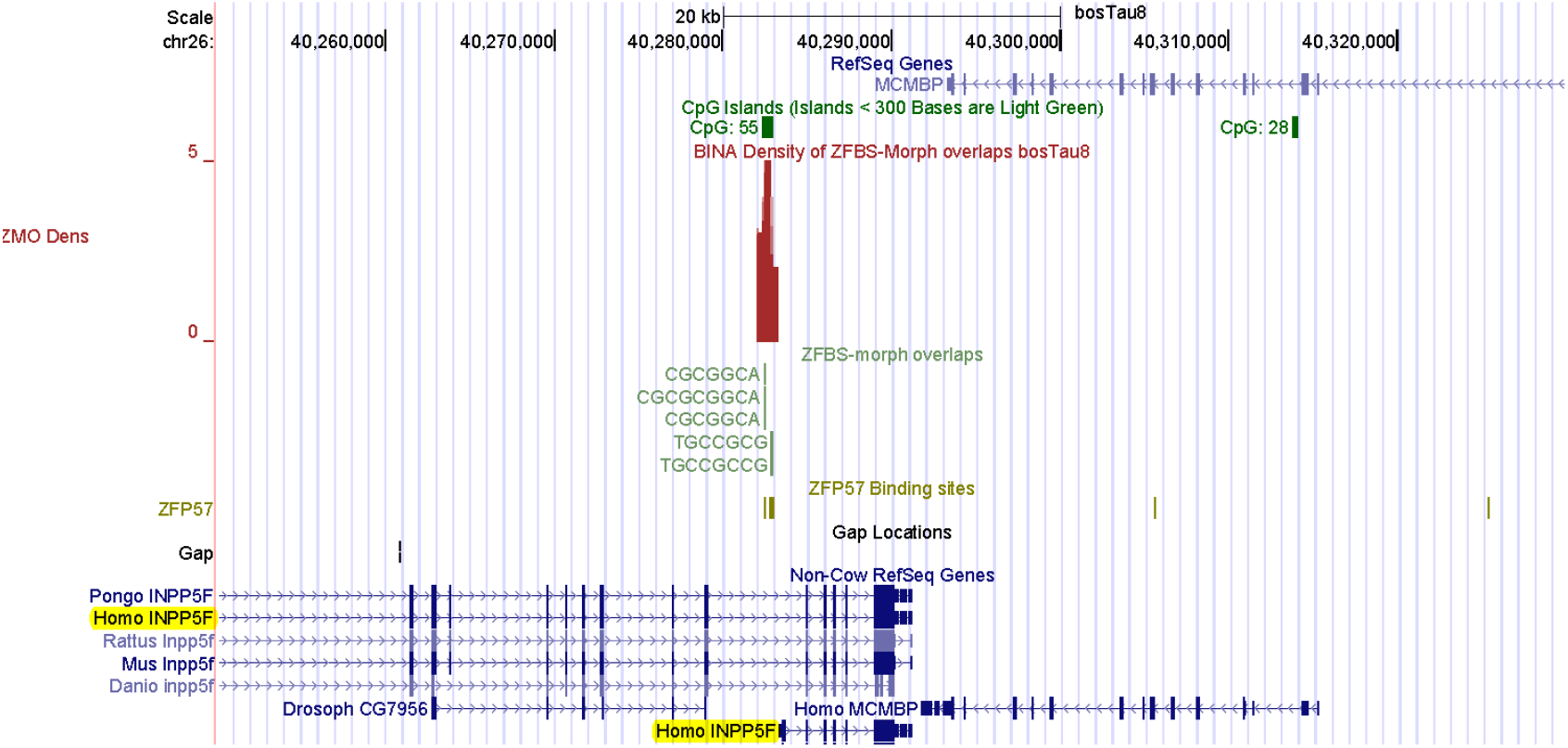
With respect to Non-Cow RefSeq Genes, a robust-peak predicted the ICR of *INPP5F_V2* in cattle DNA. As observed for mice and humans, this peak is in an intragenic CpG island.

### Density-plots could facilitate locating candidate ICRs in bovine DNA

Since the density-plots were created genome-wide, we could view peak positions along the DNA in an entire chromosome. For example, examine a snapshot from the UCSC genome browser presenting peak positions along the entire DNA in Chr26 (Fig. 11). In the build bosTau8, this chromosome covers greater than 51,7 Mb DNA. We find that even along an entire chromosome, many peaks are almost fully resolved. This plot demonstrates clearly that density peaks occur infrequently in cattle DNA. This finding is expected since CpG frequency is relatively low in animal DNA [84]. Since the peaks encompass several CpGs, they represent uncommon events along genomic DNA. Additionally, as one would expect, peaks covering 3 or more ZFBS-morph overlaps are sparser than those that covering 2 (Fig. 11). Since in plots, several peaks have correctly identified regions that include known ICRs and DMRs (Figs. 1–10), we propose that additional peaks correspond to candidate ICRs. Examples include a robust peak in a bidirectional promotor regulating *IPMK* and *CISD1*, a peak associated with *PTEN*, and two intragenic peaks regulating *SUFU*, and *JAKMIP3* expression (Fig. 11).

**Figure 11.**
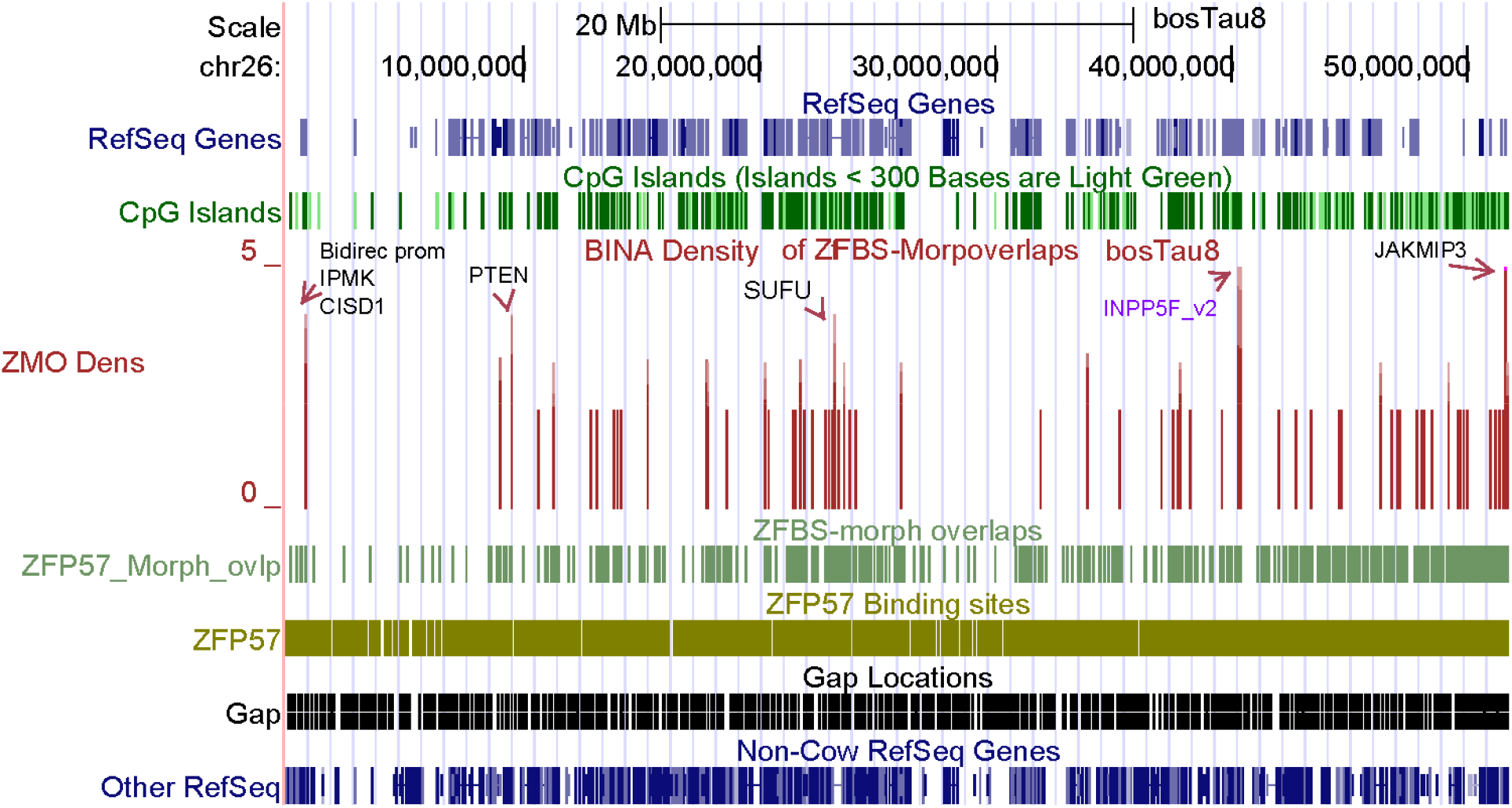
A snapshot of the density-plot obtained for the entire Chr26. Peaks covering 2 ZFBS-morph could be true or false-positives. Robust peaks cover 3 or more overlaps. Purple indicates the position of candidate ICR for imprinted *INPPF5_V2* expression. Additional robust peaks correspond to predicted ICRs. Arrows mark representative examples of such ICRs with respect to nearby genes.

To facilitate data interpretation, we have tailored our datasets for upload onto the UCSC the genome browser to create custom tracks. The browser offers a great resource for studies of genomic DNA sequences in higher organisms [85–87]. On the browser, default tracks facilitate viewing our datasets in the context of genomic landmarks including genes, transcripts, the CpG islands, SNPs and much more [88]. At the Purdue University Research Repository (PURR), you can access and download the datasets listed below:

-The positions of ZFBS and ZFBS-Morph overlaps in the build bosTau8 of the cow genome https://purr.purdue.edu/publications/3359/1
-Density of ZFBS-Morph overlaps in the build bosTau8 of the cow genome https://purr.purdue.edu/publications/3360/1

At PURR, you can also access and download datasets for studies of genomic imprinting in human, mouse, and dog. For the Bos Taurus genome, you can view peak positions along an entire chromosome (Fig. 11). You can even zoom into a desired region to obtain closeup views (Figs. 1–10). In closeup views, you can inspect peak positions with respect to genes, transcripts, CpG islands, RefSeq Genes, Non-Cow RefSeq Genes, and SNPs [87, 89]. Furthermore, it is well known that most ICRs regulate allele-specific expression of nearby genes. Therefore, since the browser is user-friendly [87], with our datasets, you can easily discover candidate ICRs and imprinted genes for experimental validations. For an overview about how to use the UCSC genome browser, see [86, 88].

## METHODS

### Marking the genomic positions of ZFP57 binding site and the ZFBS-morph overlaps

At the UCSC genome browser, we selected the build bosTau8 to obtain and download the nucleotide sequences of cow chromosomes. Initially, we wrote a Perl script to determine the genomic positions of the reported hexameric ZFP57 binding site [14], and sequences of the ZFBS-morph overlaps [48]. The script opened the file containing the nucleotide sequence of a specified chromosome, as well as either the file containing the ZFP57 binding site or sequences of the ZFBS-morph overlaps. With UNIX subroutines, we combined the outputs obtained for various chromosomes for display at the browser.

### Creating plots of the density of ZFBS-morph overlaps in genomic DNA

With a Perl script, we established the genomic positions of DNA segments that covered 2 or more closely spaced ZFBS-morph overlaps. That script opened the file containing the positions of ZFBS-morph overlaps for a specified chromosome. Subsequently, the script scanned the file to count and report the number of ZFBS-morph overlaps within a sliding window consisting of 850-bases. To remove background noise, the script ignored isolated overlaps. Next, using a subroutine we combined and tailored the outputs of the program for display as a custom track on the UCSC genome browser.

